# Herring roe oil exerts anti-psoriatic and immunomodulatory effects on the IL-17/23 signaling axis in macrophages, T-cells and a keratinocyte and fibroblast co-culture

**DOI:** 10.1101/2025.04.22.650092

**Authors:** Jennifer Mildenberger, Nina Therese Solberg, Tone-Kari K. Østbye, Vibeke Høst, Federico Petrucelli, Runhild Gammelsæter, Sissel B. Rønning, Mona E. Pedersen

**Author notes:** **Corresponding author:** Name: Jennifer Mildenberger, Address: Borgundveien 340, 6009 Ålesund, Norway. **Ethical Approval:** Isolation of primary immune cells from peripheral blood was under approval by the Regional Ethics Committee (REK, ref.: 230804). **Ethics statement:** Not applicable. **Data availability statement:** All information to reproduce the here presented analyses is included in the manuscript and supplemental information. **Author contribution:** J. Mildenberger: Conceptualization, Formal analysis, Funding acquisition, Investigation, Methodology, Project administration, Visualization, Writing – original draft, Writing – review & editing, N. Solberg: Formal analysis, Investigation, T.-K. K. Østbye: Conceptualization, Formal analysis, Funding acquisition, Investigation, Methodology, Visualization, V. Høst: Formal analysis, Investigation, F. Petrucelli: Investigation, R. Gammelsæter: Conceptualization, Funding acquisition, Methodology, Project administration, Writing – original draft, Writing – review & editing, S. B. Rønning: Conceptualization, Formal analysis, Funding acquisition, Investigation, Methodology, Visualization, Writing – original draft, Writing – review & editing, M. E. Pedersen: Conceptualization, Funding acquisition, Methodology, Project administration, Visualization, Writing – original draft, Writing – review & editing.

## Abstract

**Background:** Several systemic treatment options are available for moderate to severe psoriasis, while low-risk oral treatments for milder forms are limited. Herring roe oil (HRO) has previously shown beneficial effects on the clinical presentation of psoriasis, however, cell-specific effects remained unknown.

**Objective:** This study aimed to elucidate cellular effects of HRO and whether signaling pathways in skin and immune cells relevant to the pathophysiology of psoriasis are affected by HRO.

**Methods:** We used cell-culture models of human primary macrophages and T-cells, the keratinocyte cell line HaCat and human primary fibroblasts to represent important site-specific pathological aspects of psoriasis in the skin and immune system. The cells were supplemented with HRO emulsions and cellular signaling was analyzed on protein and mRNA level. Skin cell proliferation was analyzed in real-time by a Live-Cell Analysis System.

**Conclusion:** We here present results which suggest that HRO exerts effects on different cell types on major psoriasis-driving aspects, reducing the secretion of IL-23 and IL-17 from macrophages and T-cells, respectively, as well as interfering with IL-17-induced signaling and proliferation of keratinocytes in co-culture with fibroblasts.

Clinicaltrials.gov (or equivalent) listing (if applicable): not applicable

**Key Points:**

- Why was the study undertaken? To elucidate cellular targets for the anti-psoriatic effects of HRO.
- What does this study add? This study adds an understanding of cell specific effects of HRO that support a clinically beneficial action on psoriasis.
- What are the implications of this study for disease understanding and/or clinical care? This study supports the use of HRO as treatment option for mild-to-moderate psoriasis.

## Introduction

Psoriasis is a chronic immune-mediated inflammatory disease (IMID) affecting the skin. The exact etiology is unknown but involves genetic and environmental influences. The physical manifestations of psoriasis are characterized by hyperproliferation and abnormal differentiation of keratinocytes, leading to scaly plaques, and by a strong inflammatory component manifesting in massive infiltration of immune cells^1,2^. It is recognized that the regulation of skin immune functions happens in the epithelial immune microenvironment between keratinocytes in the epidermis and immune cells in the papillary dermis^3,4^.

In the development of psoriasis, keratinocytes are triggered by external stimuli to release cytokines and antimicrobial peptides (AMPs). These form a complex with nucleic acids and are sensed by plasmacytoid dendritic cells (pDC) and macrophages leading to the secretion of Interleukin 1-alpha (IL-1α) and Tumor necrosis factor alpha (TNF-α), further stimulating the maturation of resident dermal DCs and infiltrating monocyte-derived inflammatory DCs^5^. Mature and activated DCs secrete cytokines such as IL-12, IL-23, Interferon gamma (IFN-γ) and TNF-α and present antigens to naïve T-cells, which differentiate into Th1, Th17 and Th22 helper cells. Th1 and Th17 cells secrete mainly IFN-γ and IL-17, respectively^6^. The secretion of Th17 cytokines promotes keratinocyte hyperproliferation and cytokine secretion, again stimulating T-cells and attracting DCs, neutrophils and macrophages^7–9^. Despite much focus on DCs, macrophages are involved in both initiation and sustained inflammation in psoriasis. They shift to a pro-inflammatory polarization characterized by secretion of IL-23, TNF-α and IL-1β, as well as neutrophil-attracting chemokines, all with a driving role in psoriasis^3^. Psoriatic patients have increased numbers of circulatory monocytes and in mouse models, depletion of macrophages has ameliorated clinical signs of psoriasis^5,10^.

Both keratinocytes and fibroblasts are affected by and contribute to the aberrant inflammatory signaling in the psoriatic skin. Keratinocytes express IL-17 receptor and binding of IL-17 activates the transcription factor nuclear factor kappa-light-chain-enhancer of activated B cells (NF-κB) and mitogen-activated protein kinases^7^. The NF-κB inhibitor zeta (*NFKBIZ*, encoding IκBζ protein) has recently been shown to be a positive regulator of NF-κB and central mediator of IL-17-driven effects^11–13^. IκBζ in keratinocytes is further involved in the upregulation of psoriasis-associated genes, such as the AMP and psoriasis marker S100A7 (also known as psoriasin)^14–16^. S100A7 has been linked to keratinocyte hyperproliferation and impaired differentiation and overall amplified inflammation^17,18^. Fibroblasts are also targets of IL-17 and capable of sustaining inflammation by the production of proinflammatory mediators and growth factors^19,20^. Notably, fibroblasts are one of the first cell types to respond to blockage of IL-23^21^.

In moderate-to-severe psoriasis, biologics with effect on the IL-17/IL-23 axis have shown successful treatment outcomes, interfering with the vicious cycle of over-activation^1,22^. However, there is a lack of well-tolerated oral treatment options for mild-to-moderate psoriasis.

Omega-3 polyunsaturated fatty acids (n-3 PUFAs) have established immunomodulatory effects, which are relevant to the amelioration of psoriasis^23–25^. They are further metabolized to specialized pro-resolving mediators (SPMs)^26^ with a therapeutic potential in IMIDs^27^. Nevertheless, there is no consensus of anti-psoriatic effects of n-3 PUFAs in clinical trials^28^. Herring (*Clupea harengus*) roe oil (HRO) has a high content of phospholipid esters enriched with marine n-3 PUFAs, such as eicosapentaenoic acid (EPA) and docosahexaenoic acid (DHA). HRO has previously been shown to improve the clinical manifestation in patients with mild *Psoriasis vulgaris*^29,30^, which was supported by immunomodulatory effects^31^. Supplementation of HRO in macrophages and keratinocytes has also led to the biosynthesis of SPMs^32^. We have here characterized effects of HRO on cellular responses of immune and skin cells that are relevant in the context of psoriasis to contribute to the elucidation of the mode-of-action of HRO.

## Material and methods

### Herring roe oil

Herring (*Clupea harengus*) roe was used for extraction of HRO as previously described^32^. A 5 % (w/w) emulsion in dH_2_O was prepared by stirring at 800 rpm under inert atmosphere (nitrogen flushing) at 37°C for 1 h. Emulsions were then pasteurized under inert atmosphere by sonication at 70°C for 2 min in an ultrasonic water bath (Branson 2200, 40 kHz) or by an ultrasonic probe. Emulsions were stored at 4°C for maximum 4 days prior to cellular experiments.

### Cell culture

Buffy coats were obtained from the blood bank Ålesund (Helse Møre og Romsdal, Norway) with approval by the Regional Ethics Committee (REK, ref.: 230804). Peripheral blood mononuclear cells (PBMC) were isolated from buffy coats by density gradient and monocytes were selected by plastic adherence and differentiated into monocyte-derived macrophages (MDM) by addition of macrophage colony-stimulating factor (M-CSF). To induce inflammatory signaling, cells were treated with 1 µg/mL lipopolysaccharide (LPS, Sigma-Aldrich, L2630) and 50 ng/mL IFN-γ (Miltenyi Biotec, 130-096-481).

CD4+ T-cells were isolated from PBMC by negative magnetic bead separation using the CD4^+^ T Cell Isolation Kit (Miltenyi Biotec, 130-096-533). T-cell purity was checked and was routinely above 95 % (example in Supplemental Figure 1). T-cells were activated by CD3/CD28 Dynabeads™, at 1 bead/2 cells (Gibco, 11161D) and cytokines (IL-1β (20 ng/mL, Invitrogen, A42508), Prostaglandin E2 (PGE_2_) (5 µM, Cayman chemicals, 14010), IL-23 (25 ng/mL, Miltenyi Biotec, 130-095-757) for 3 days.

THP1-Dual™ NF-κB-SEAP IRF-Luc Reporter monocytes (InvivoGen, thpd-nfis) were grown as recommended by the manufacturer.

Human primary dermal fibroblasts (ATCC PCS-201-012) were cultured for 2 days before the addition of human keratinocytes HaCaT (CLS Cell line service GmbH, cat no 300493), either in a separate well or to the same well as fibroblasts for direct co-culture, or the filter for co-culture in Transwell® (TW) plates (Corning Costar Transwell, VWR). For TW-co-culture experiments, fibroblasts were seeded at well bottom. Cells were further co-cultured for 5 days before stimulation with or without 100 ng/mL IL-17A for 4 days (Sigma-Aldrich, cat.no SRP0675). Real-time cell confluence was monitored every 4 h using an IncuCyte® S3 Live-Cell Analysis System (Sartorius AG, Germany).

All cells were maintained in 5 % CO_2_ at 37°C and full cell culture details are provided in the supplemental data (Appendix⍰S1, Supporting Information).

### Reporter assays

NF-kB or IRF activation were analyzed by secreted embryonic alkaline phosphatase (SEAP) (QUANTI-Blue™, InvivoGen, rep-qbs) or Lucia luciferase (QUANTI-Luc™, InvivoGen, rep-qlc1) reporter assay in cell supernatants from THP1 reporter cells after stimulation with 10 ng/mL LPS for 24 h, according to the manufacturer’s instructions. QUANTI-Luc™ was analyzed immediately by luminescence and QUANTI-Blue™ after 30 – 60 min of incubation by absorbance at 635nm on a Synergy HTX S1LFA plate reader (BioTek).

### Enzyme-linked immunosorbent assay (ELISA)

Cytokine secretions were analyzed in duplicates in appropriately diluted medium from treated cells by human IL-23 ELISA kit (Invitrogen, 88-7237-88) or Duo ELISA kits (R&D technologies) for human CXCL10 (DY266), IL-17A (DY317), TNF-α (DY 210), IFN-γ (DY285B) and IL-10 (DY217B) according to the manufacturer’s instructions. Absorbance was analyzed on a Synergy HTX S1LFA reader (BioTek). GainData®^,33^ was used with a 4-parameter logistic regression standard curve for the calculation of cytokine concentrations.

### Quantitative real-time PCR (qrt-PCR)

RNA was isolated by spin-columns, cDNA was generated by reverse transcription, and qrt-PCR was run with primer- and probe-based detection chemistries. Relative mRNA levels were transformed into linear form by the 2 (−ΔΔCt) method^34^. Detailed protocols are provided in the supplemental data (Appendix⍰S1, Supporting Information).

### Western Blot

Protein was isolated by lysis in RIPA buffer and separated on SDS-PAGE NuPage 12 % Bis-Tris gels (Invitrogen), before transferred to nitrocellulose membranes (iBlot3 Transfer stacks, IB33001, Invitrogen) using an iBlot Gel Transfer system (Invitrogen). All membranes were blocked with 2 % ECL Prime blocking agent (RPN418V, Cytiva), before incubating with primary and secondary antibodies. Detailed protocols are provided in the supplemental data (Appendix1S1, Supporting Information).

### Statistical analysis

Statistical analyses were performed in GraphPad Prism 10.0.1. Data of primary immune cells was analyzed by non-parametric Friedmans test, data from all other cells was analyzed by ANOVA. Shown are boxplots from min. to max. showing all points or bar graphs for means with standard deviation. P-values of <0.05 were considered significant and only statistically significant comparisons are shown for all figures.

## Results

As a first step for the characterization of the cellular mode-of-action of HRO, the range of tolerable supplementation was analyzed. Primary MDMs, THP-1 macrophages and T-cells were treated with HRO for 24 h and keratinocytes and fibroblasts were treated for 96 h, according to the further treatment set-up. Up to 500 µg/mL HRO were tolerated in the immune cells and 50 µg/mL HRO in the skin cells without impairment of cellular viability (Supplemental Figure 2). Cellular uptake of fatty acids from HRO has been described before, reporting significant increases in the n3-PUFAs EPA and DHA in the here used MDMs and keratinocyte/fibroblast co-culture^32^.

### HRO supplementation limits secretion of IL-23 from macrophages

On the cellular level, the development and progression of psoriasis is driven by the reciprocal over-activation of immune and skin cells^35,36^. Macrophages and DCs in the skin produce, among other cytokines, IL-23 as an important driver of the inflammatory cycle in psoriasis^37,38^. Macrophages were selected for the evaluation of HRO effects, due to their more consistent induction of IL-23 (data not shown) and previously described use as models for IL-23 signaling^39^. MDMs were pretreated by HRO overnight, followed by stimulation with LPS and IFN-γ. Secretion of IL-23, TNF-α and CXCL10 were all significantly reduced by 500 µg/mL HRO and IL-23 also by 50 µg/mL HRO (Figure 1a-c). On mRNA level, the expression of *IL23* was similarly reduced after pretreatment with 500 µg/mL HRO, as well as the expression of *TNFA* and *IFNB* as targets of NF-κB or IRF, respectively (Figure 1d-f). In a set-up with short inflammatory stimulation (2 h), HRO was effective when THP-1 cells were pretreated with HRO for 16 h, but not when pretreated for 2 h or HRO added at inflammation induction (Supplemental Figure 3). Finally, we observed a strong reduction of both NF-κB and IRF activity in THP-1 reporter cells, suggesting that dampening effects of HRO occur upstream of transcription factor activation and on several pathways (Figure 1g,h).

**Figure 1.**
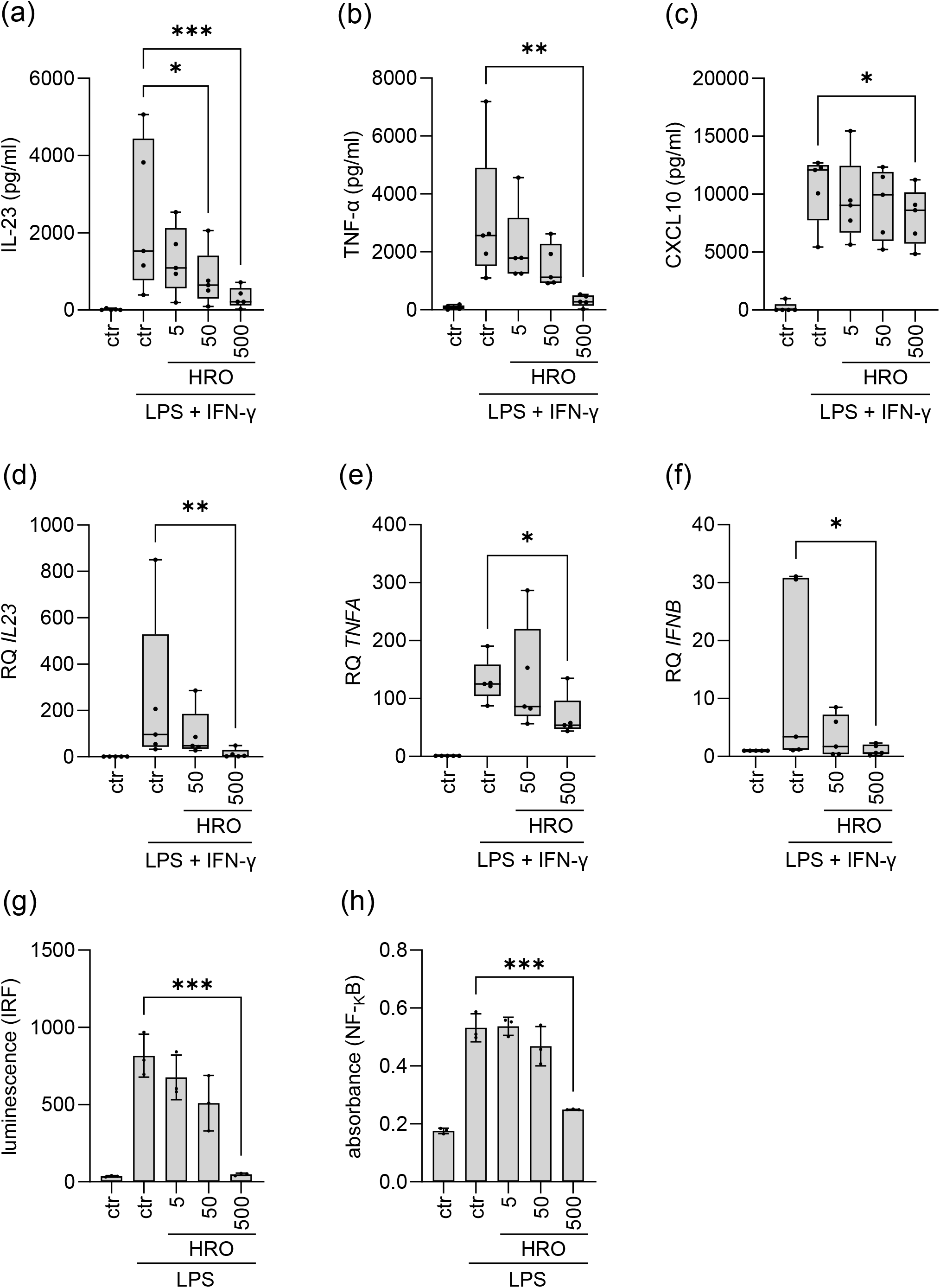
Human primary MDMs were pretreated with HRO at indicated concentrations (5 - 500 µg/mL) for 16 h, followed by stimulation with LPS (1 µg/mL) and IFN-γ (50ng/mL) for 24 h and analysis of secreted IL-23 (a), TNF-α (b) and CXCL10 (c) by ELISA (n=5). Or followed by stimulation with LPS and IFN-γ for 4 h for assessment of mRNA levels by qrt-PCR for IL23 (d), TNFA (e), and IFNB (f) (n=5). Reporter activity of NF-κB (g) and IRF (h) reporters in THP-1 macrophages treated with HRO as indicated (5 -500 µg/ml) for 16 h, followed by stimulation with 10 ng/ml LPS for 24 h (n=3). All treatments were compared to the LPS-stimulated controls. * = P ≤ 0.05; ** = P ≤ 0.01; *** = P ≤ 0.001

### HRO supplementation limits IL-17 secretion from T-cells

Increased levels of IL-23 lead to the differentiation of naive T-helper-cells into Th17-cells, which secrete high amounts of IL-17^6^. T-cells were treated with combinations of IL-1β, PGE_2_ and IL-23 for 3 days to induce secretion of IL-17^40^. Co-treatment with HRO did significantly reduce IL-17 secretion for all inducing stimulations (Figure 2a). TNF-α was not induced by stimulation but still reduced by HRO (Figure 2b). IFN-γ was induced by stimulation and reduced by HRO, suggesting that also Th1-cells might be affected by HRO supplementation (Figure 2c). IL-10 was suppressed by stimulation with PGE_2_ as described before^40^ and not further changed by HRO (Figure 2d). Complementary analysis by Flow cytometry for Th17 surface markers was not successful due to the needed prolonged time for differentiation into Th17-cells, which was accompanied by degradation of supplemented HRO. IL-17 was also induced by co-culture of T-cells with activated MDMs as described before^39^. IL-17 secretion in this model was significantly reduced when either T-cells or the co-culture was supplemented with HRO (Figure 2e), while pretreatment of MDMs was not effective.

**Figure 2.**
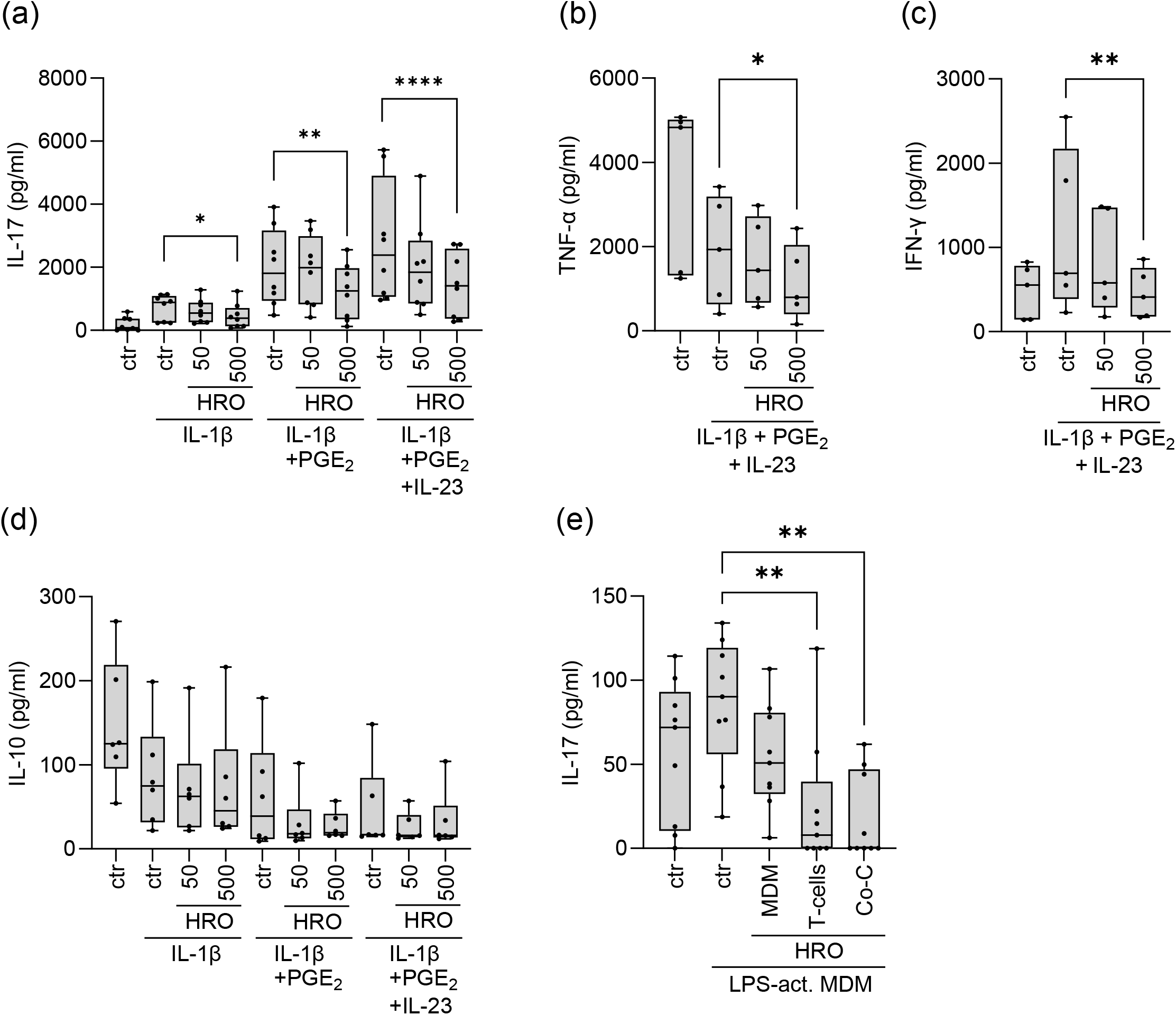
IL-17 (a), TNF-α (b), IFN-y (c) and IL-10 (d) secretion assessed by ELISA from activated T-cells stimulated with indicated combinations of IL-1β (20 ng/ml), PGE_2_ (5µM) and IL-23 (25 ng/ml) in presence or absence of 50 or 500 µg/ml HRO for 3 days (n*≥*5). (e) IL-17 secretion assessed by ELISA from co-cultures of activated MDMs and autologous T-cells. Indicated cell types were treated by HRO (50 µg/ml). MDMs were pretreated by HRO for 16 h, then activated (LPS-act. MDM) by LPS (1µg/ml) for 24 h, followed by addition of T-cells. T-cells were pretreated with HRO for 16 h before addition to MDMs. Addition of HRO upon initiation of the co-culture is indicated as Co-C (n=9). HRO treatments were compared to the corresponding stimulated control without HRO treatment. * = P ≤ 0.05; ** = P ≤ 0.01; **** = P ≤ 0.0001

### HRO reduces the expression of NFKBIZ and S100A7, and proliferation of co-cultured keratinocytes and fibroblasts

IL-17 activates keratinocytes in the skin, increasing their cytokine secretion, and interfering with proliferation and differentiation^2,7–9^. HRO did not change the IL-17-stimulated secretion of inflammatory cytokines from keratinocytes in TW-co-culture with fibroblasts (Supplemental Figure 4). However, in a physical co-culture of keratinocytes and fibroblasts, HRO significantly reduced the IL-17-induced expression of both *NFKBIZ* and its downstream target *S100A7* (Figure 3a,b), while at the protein level the reduction in S100A7 was non-significant across biological replicates (Figure 3c). We further analyzed the proliferation of co-cultured keratinocytes and fibroblasts that was not significantly affected by neither IL-17 nor 5 µg/mL HRO, or their combination (Figure 3d). However, with 50 µg/mL HRO we observed a significant reduction in proliferation after 70 hours, both with and without IL-17. Investigations of keratinocytes and fibroblasts when co-cultured in a TW-system, or monocultured for proliferation studies, revealed that both cell types reduced the IL-17-induced expression of *NFKBIZ* with 5 µg/mL HRO (Supplemental Figure 5a,d). Similar to the observations in the physical co-culture, the IL-17-induced expression of S100A7 was reduced by addition of 5 µg/mL HRO in keratinocytes, and proliferation was reduced with 50 µg/mL HRO, regardless of IL-17 treatment (Supplemental Figure 5b,c). However, in fibroblasts 5 µg/mL HRO significantly enhanced both the IL-17-mediated expression of S100A7, and significantly increased proliferation after 30 hours, regardless of IL-17 stimulation (Supplemental Figure 5e,f). These data identify opposite effects of HRO on keratinocytes and fibroblasts, with respectively reduced and increased proliferation. 50 µg/mL HRO resulted in a cloudy cell media, over-estimating the confluency measurements of both cell types (data not shown). This effect was visually reduced over time by keratinocytes, both alone and when co-cultured with fibroblasts, but not by fibroblast monoculture. This observation explains the bumpy 50 µg/mL HRO graphs in Figure 4d and Supplementary Figure 4c as well as the high confluency graphs in Supplemental Figure 5f.

**Figure 3.**
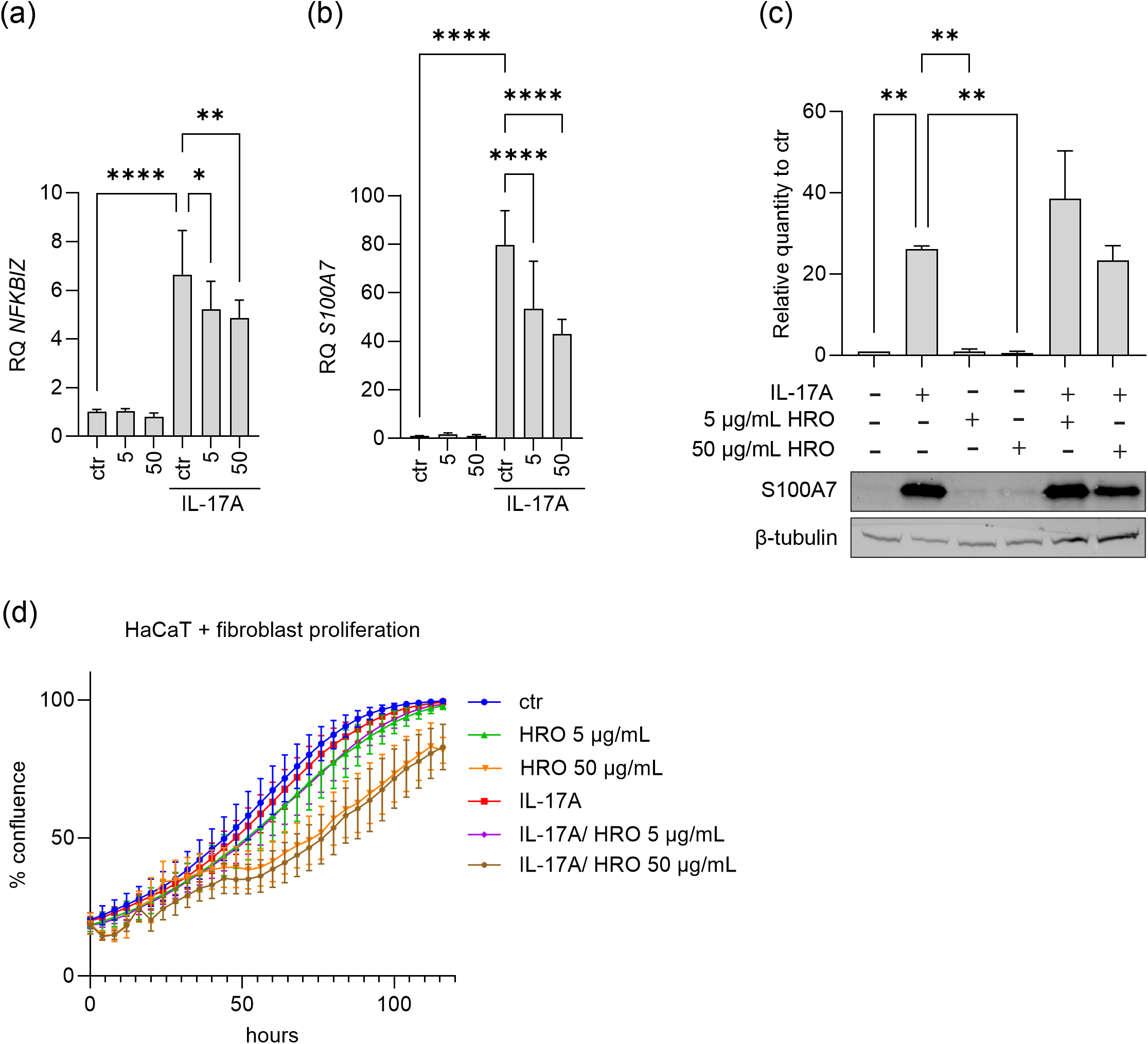
RT-qPCR analysis of the psoriasis related response genes *NFKBIZ* (a) and *S100A7* (b) (n=3), and Western blot analysis of the S100A7 protein (c) (n=3) on RNA and protein isolated from a physical co-culture of keratinocytes and fibroblasts. Keratinocytes were co-cultured with fibroblasts for 5 days before stimulation with 100 ng/mL IL-17A, and/or 5 or 50 µg/mL HRO for 4 days. For proliferation analysis keratinocytes and fibroblasts were co-cultured in a 5:1 ratio for 1 day before 100 ng/mL IL-17A, and/or 5 or 50 µg/mL HRO was added (d). Monitoring of cell confluence was started immediately after addition of stimuli and subsequently monitored every 4 hours for 120 hours. The proliferation curve shown is a representative of 4 graphs, each with 4 technical replicates. Western blot analysis was performed on 3 biological replicates, each with 1 technical replicate. Representative picture is shown. All treatments were compared to each other. For RT-qPCR: only statistical significance to control (ctr) with IL-17A stimuli are shown. * = ≤ 0.05; ** = P ≤ 0.01; **** = P ≤ 0.0001.

## Discussion

Psoriasis has a complex pathophysiology including the inflammatory interaction of many skin and immune cell types^41^. While there is a plethora of systemic treatments indicated for patients with moderate-to-severe psoriasis, there are limited treatment options for mild-to moderate psoriasis. Patients with non-severe psoriasis may not have marked systemic inflammation, and immunosuppressive treatments may not be appropriate. HRO has previously shown beneficial effects on clinical signs in psoriasis^29,30^, and decreased activation of neutral killer cells and monocytes systemically^31^. Here, we have characterized effects of HRO on models of skin and immune cells present in psoriatic skin and demonstrated anti-psoriatic and immunomodulatory effects. HRO decreased the secretion of IL-23, TNF-α and CXCL10 from MDMs, further in line with their decreased expression and limited activation of NF-κB and IRF. The impact of HRO on the activation of these central inflammatory transcription factors suggests that also other downstream cytokines might be reduced by HRO. N-3 PUFAs are well known to exert anti-inflammatory effects via multiple pathways^25,42^ that could be further investigated for HRO. In case of short inflammatory stimulation (2 h) as for expression analysis of cytokines, HRO was only effective as pretreatment. This suggests that time for intracellular signaling or metabolism is needed for HRO to exert its observed anti-inflammatory effect. HRO-supplemented MDMs have been shown to produce SPMs^32^ with established resolving actions^27^ and a causative connection to the here presented results should be investigated in future studies. Likewise, the SPM Resolvin D1 (RvD1) has been shown to limit NF-κB activation and cytokines of the IL23/IL17 axis^43^ and RvE3 reduced IL-23 release from macrophages^44^. HRO supplementation also limited IL-17 and IFN-γ secretion from T-cells, thus markedly interfering with inflammatory T-cell signaling. Also, pretreatment of T-cells with HRO led to reduced IL-17 secretion when induced by co-culture with MDMs. The ineffective HRO pretreatment of MDMs in this case might have been due to the sequence of the experiment, where HRO treatment of T-cells or the co-culture was done more shortly before analysis. This *in vitro* set-up has also limitations as the chosen supplementation regime for HRO could not be sustained during enough time for complete differentiation into Th17 cells and should be optimized further. However, n-3 PUFAs as well as RvD1 have previously been shown to reduce Th17 differentiation in other models^45–48^. Hence, HRO interferes with the inflammatory circle in psoriasis by dampening both IL-23, TNF-α and IL-17 directly from secreting immune cells. These are the main cytokines also targeted by biologics for the treatment of more severe psoriasis, underlining the clinical relevance of HRO-mediated effects^1,22^. We did not observe effects of HRO on IL-17-induced cytokine secretion from keratinocytes, suggesting a differentiated action of HRO on different signaling pathways, unlike an immunosuppressive mechanism. HRO reduced the IL-17-induced expression of *NFKBIZ* and its downstream target *S100A7*. IκBζ is an important driver of IL-17 mediated psoriatic effects and an interesting therapeutic target^13^. In the fibroblast compartment of the TW, *NFKBIZ* was reduced while *S100A7* was increased by HRO, suggesting a differential regulation in the epidermis and dermis. The non-significant effect of HRO on S100A7 protein level measured in direct co-culture is in line with the opposite effects on *S100A7* expression observed in these cell types. We also observed a dampening effect of HRO on keratinocyte proliferation that corresponds to the reduction of scaling in psoriatic skin in a clinical trial^30^. The anti-proliferative effect of HRO was observed independently of IL-17, which might indicate a restoring activity of HRO in case of inflammatory challenge, but also a regulatory role in normal conditions. S100A7 promotes proliferation^49^ in line with our data, where a reduction in its expression by HRO coincides with reduced proliferation. Stimulation of fibroblasts *in vitro* with recombinant S100A7 has anti-proliferative and anti-fibrotic effects^50^. Further studies are needed to clarify molecular mechanisms behind the effect of HRO on fibroblasts. In a psoriatic skin model, the n-3 PUFAs DHA, EPA and alpha linoleic acid (ALA) have reduced keratinocyte proliferation and DHA and ALA also normalized differentiation^51–53^. It is important to keep in mind that *in vitro* models cannot reflect the timeline *in vivo*, where it took several months before clinical effects were seen^29,30^, however, allowing the investigation of local effects that cannot be measured systemically. Overall, here we have focused on some major aspects of psoriasis pathophysiology and, as HRO has shown effects on all here tested cell types, HRO might exert even further effects on the numerous cell types and pathways involved in psoriasis.

## Conclusion

HRO displays anti-inflammatory effects on macrophage and T-cell signaling relevant for psoriasis through limiting secretion of IL-23 and IL-17, respectively. Reduction of *NFKBIZ* and *S100A7* along with reduced proliferation of keratinocytes, support an anti-psoriatic effect of HRO on skin cells.

The observed effects on the IL-23/IL-17 axis may potentiate a mechanism of action where HRO may affect several cell types simultaneously. Taken together with recent data demonstrating that HRO is taken up to these cell types and promoting broad lipid mediator biosynthesis^32^, this may indicate that HRO can impact intracellular processes associated with psoriasis in multiple cell types. The clear dampening effect on inflammation in immune cells could also be relevant as a treatment modality in other IMIDs and warrants further investigation.

## Acknowledgements

We would like to thank Thomas Ringheim-Bakka, Maftuna Busygina and Daniele Mancinelli from Arctic Bioscience AS who contributed to study conceptualization and provision of HRO used in the investigations. We also thank Bodil Stige from Medisinsk biokjemi og blodbank Ålesund, Helse Møre og Romsdal HF for the provision of buffy coats and Yanran Cao from NTNU Ålesund for help with Flow cytometry of T-cells.

## Abbreviations

ALA: Alpha linoleic acid
AMPs: Antimicrobial peptides
CXCL10: C-X-C motif chemokine ligand 10
DC: Dendritic cell
DHA: Docosahexaenoic acid
ELISA: Enzyme-linked immunosorbent assay
EPA: Eicosapentaenoic acid
HRO: Herring roe oil
IFN: Interferon
IκBζ: NF-kB inhibitor zeta (protein)
IL: Interleukin
IMID: Immune-mediated inflammatory disease
IRF: Interferon regulatory factor
LPS: Lipopolysaccharide A
M-CSF: Macrophage colony-stimulating factor
MDM: Monocyte-derived macrophages
N3-PUFA: Omega-3 polyunsaturated fatty acid
NF-kB: Nuclear factor kappa B
NFKBIZ: NF-kappa-B inhibitor zeta
PBMC: Peripheral blood mononuclear cells
PGE_2_: Prostaglandin E2
RvD1: Resolvin D1
S100A7: S100 calcium-binding protein A7 (psoriasin)
SEAP: Secreted embryonic alkaline phosphatase
SPM: Specialized pro-resolving mediator
TNF-α: Tumor necrosis factor alpha
TW: Transwell
Qrt-PCR: Quantitative real-time PCR

