## Supplemental figures for "Herring roe oil exerts anti-psoriatic and immunomodulatory effects on the IL-17/23 signaling axis in macrophages, T-cells and a keratinocyte and fibroblast co-culture"

**Supplemental Figure 1**

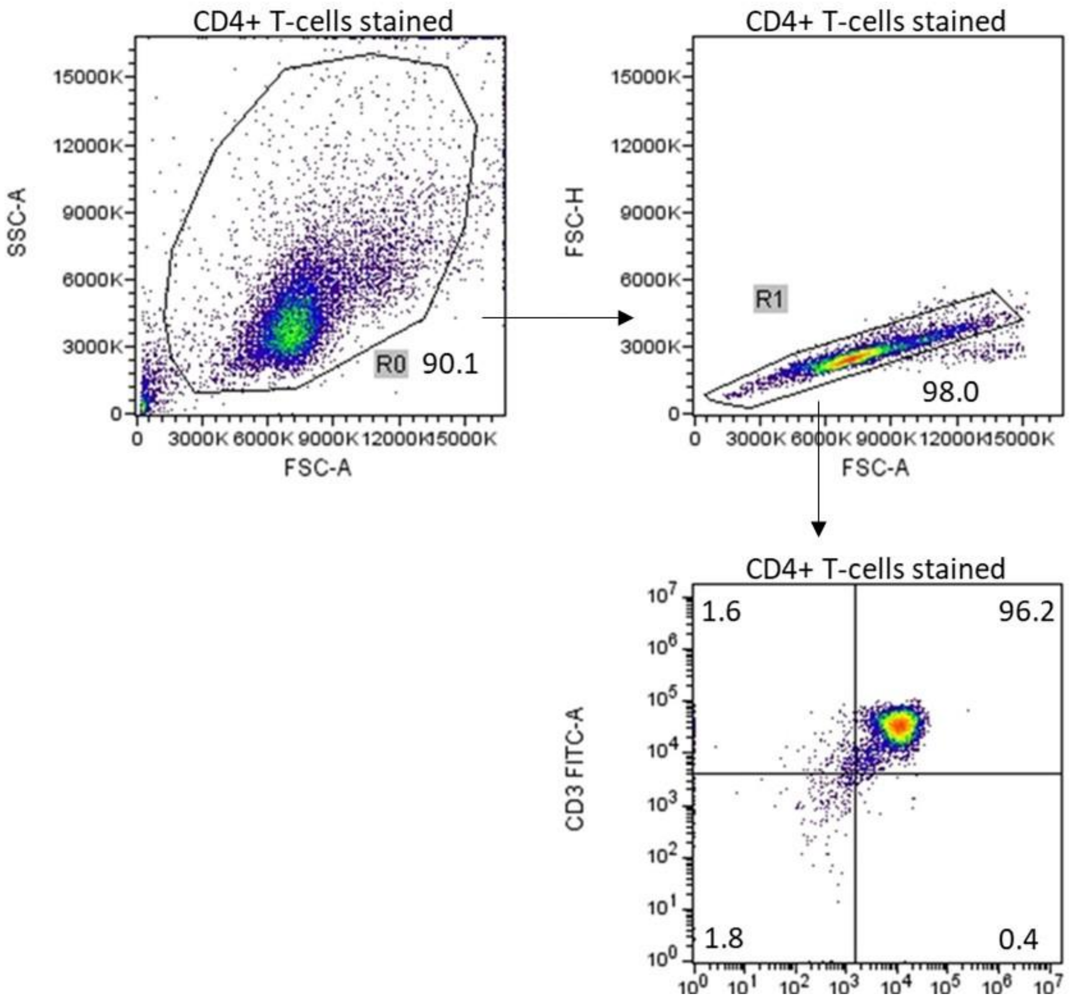


Supplemental Figure 1. Example of T-cell purity analysis and gating strategy. T-cell purity was checked by staining for CD4, CD3 and CD14, analyzed by Flow cytometry and visualized in FCSalyzer 0.9.22-alpha, and was routinely above 95 %.

**Supplemental Figure 2**

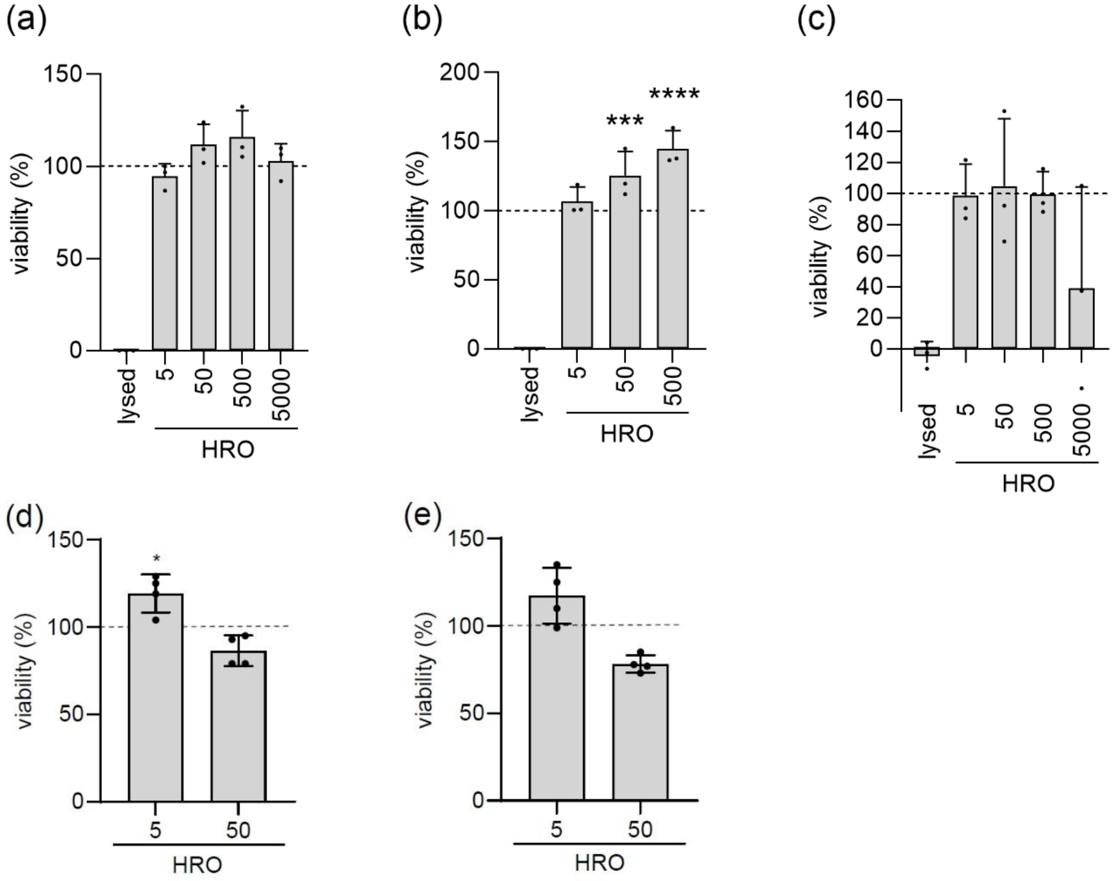


Supplemental Figure 2. Viability after treatment with indicated concentrations of HRO (5 - 5000 µg/ml) for 24 h in human primary MDMs (a), THP-1 macrophages (b) and human primary $CD4^+$ T-cells (c), and after treatment with HRO (5 and 50 µg/ml) for 96 h in HaCat keratinocytes (d) and human primary fibroblasts (e). Shown are relative viability (%) as compared to untreated control indicated as dashed line (100 %). All treatments were compared to the untreated control. * = P ≤ 0.05; *** = P ≤ 0.001; **** = P ≤ 0.0001

**Supplemental Figure 3**

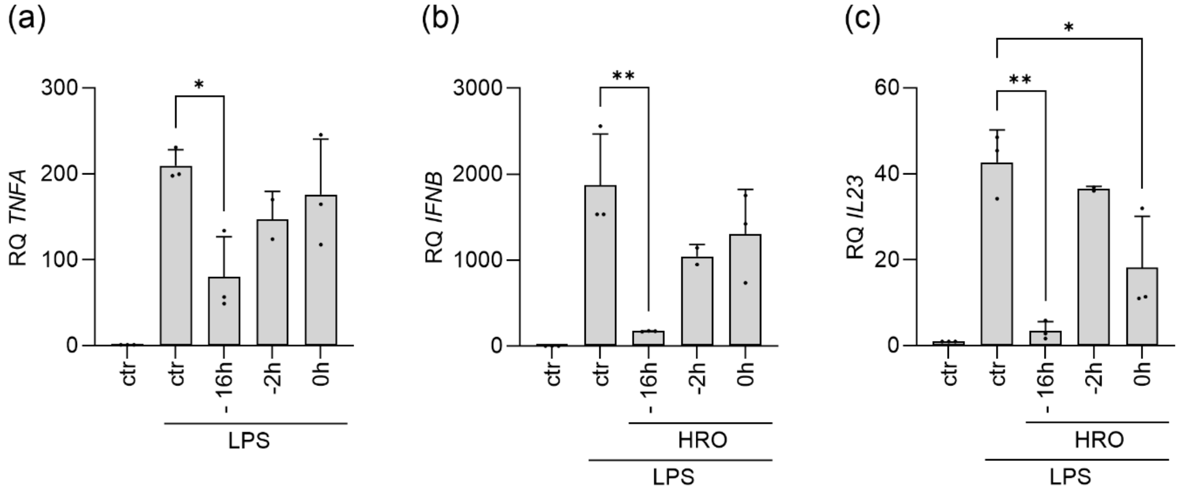


Supplemental Figure 3. Timeseries of HRO treatment (50 µg/ml) in THP-1 macrophages. The time of HRO addition (-16 h, -2 h) is indicated as time before stimulation with 10 ng/ml LPS for 2 h. Shown are relative mRNA expression (RQ) as assessed by qrt-PCR for TNFA (a), or IFNB (b) and IL23 (c). All treatments were compared to the LPS-stimulated control. * = P ≤ 0.05; ** = P ≤ 0.01

**Supplemental Figure 4**

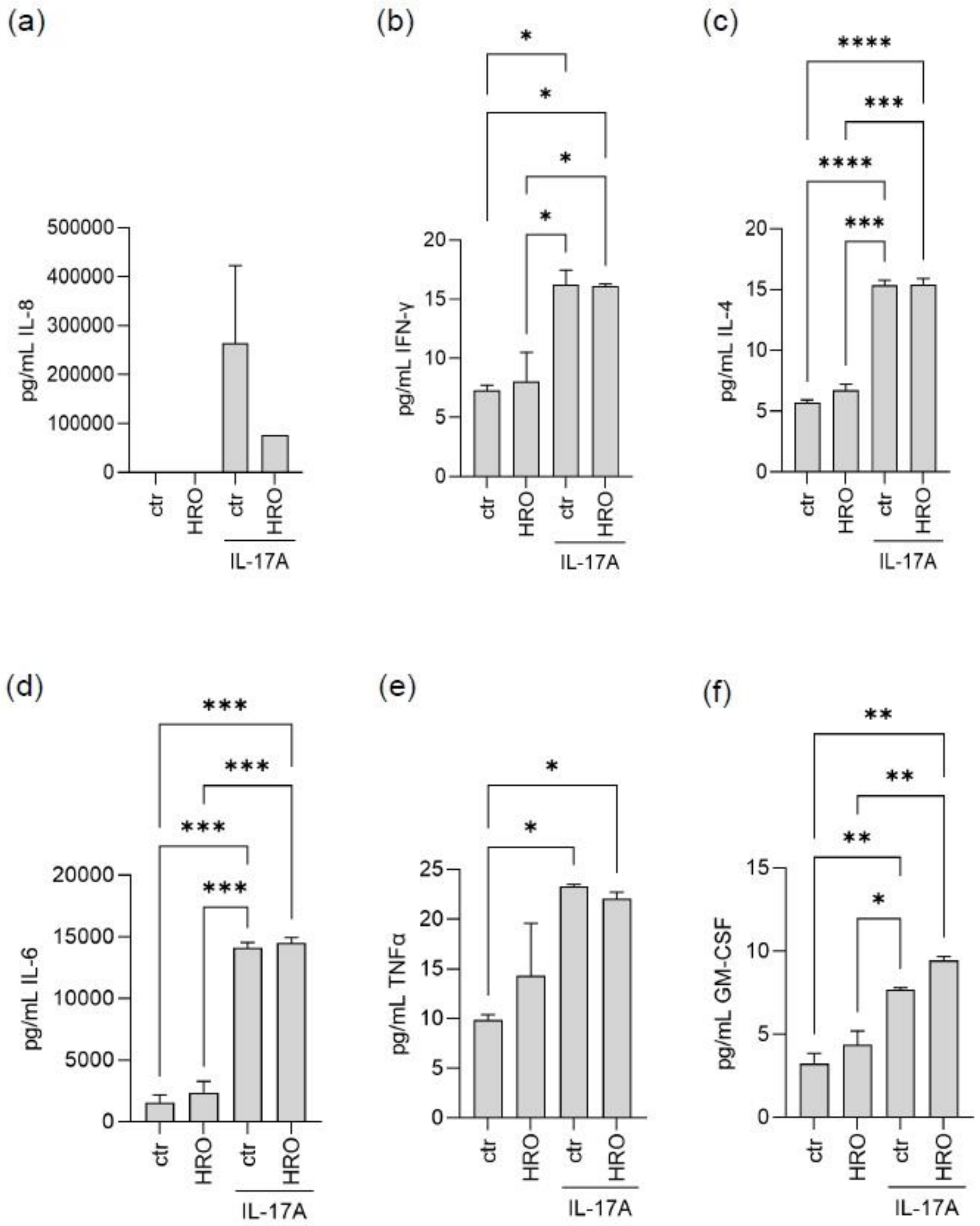


Supplemental Figure 4. IL-8 (a), IFNγ (b), IL-4 (c), IL-6 (d), TNFα (e) and GM-CSF (f) secretion assessed by BioPlex ELISA on cell media from keratinocytes co-cultured with fibroblasts in TransWells after 4 days stimulation with 100 ng/mL IL-17A and/or 5μg/mL HRO. All treatments were compared to each other. Only statistical significant comparisons are shown. * = ≤ 0.05; ** = P ≤ 0.01; *** = P ≤ 0.001; **** = P ≤ 0.0001.

**Supplemental Figure 5**

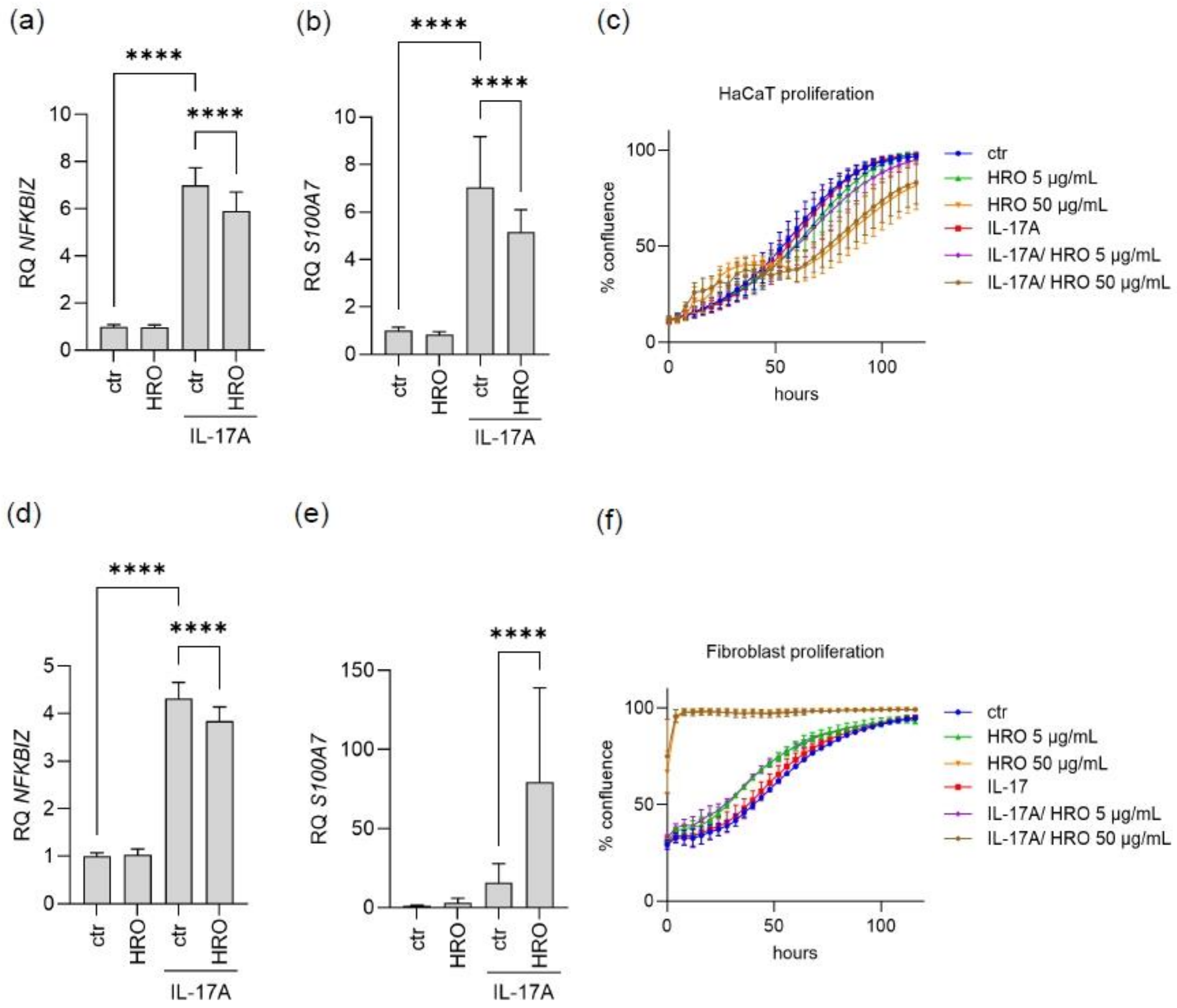


Supplemental Figure 5. RT-qPCR analysis of the psoriasis related response genes *NFKBIZ* (a) and *S100A7* (b) in mono-cultured keratinocytes, and in mono-cultured fibroblasts (d) and (e). Cell confluency of mono-cultured keratinocytes (c) and mono-cultured fibroblasts (f) plated 1 day prior to addition of 100 ng/mL IL-17A, and/or 5 or 50 µg/mL HRO. Monitoring of cell confluence was started immediately after addition of stimuli and subsequently monitored every 4 hours for 120 hours. RT-qPCR analysis was performed on 3 biological replicates, each with 3 technical replicates. The proliferation curves shown are representatives of 4 graphs each, each performed with 4 technical replicates. All treatments were compared to each other. Only statistical significant comparisons are shown. **** = $P \leq 0.0001$.
