## Appendix S1 Material and Methods for "Herring roe oil exerts anti-psoriatic and immunomodulatory effects on the IL-17/23 signaling axis in macrophages, T-cells and a keratinocyte and fibroblast co-culture"

**Cellular viability**

Effects on cellular viability of the immune cells were analyzed by a resazurin-based assay (PrestoBlue^TM^; Thermofisher, A13261) and spectrophotometric detection on a Synergy HTX S1LFA plate reader (BioTek) according to the manufacturer’s instructions. The viability of fibroblasts and HaCat was evaluated using CellTiter-Glo® Luminescent Cell Viability Assay (Promega) following the manufacturer’s protocol. Viability percentages were calculated relative to untreated control cells (100 % viability).

**RNA extraction and Real-Time quantitative PCR (RT-qPCR)**

The immune cells were lysed and RNA isolated by the Quick-RNA Miniprep Kit (Zymo Research, R1054). RNA concentration was measured by 260/280 absorbance on a Synergy HTX S1LFA plate reader (BioTek). cDNA was synthesized by the iScript cDNA kit (BioRad, 1708890) and run with the PrimePCR™ Probe Assays (BioRad) for human *TNF (qHsaCEP0040184)*, *CXCL10 (qHsaCEP0053880)*, *IFNB1 (qHsaCEP0054112)*, *IL23 (qHsaCJP0034602)*, *HPRT1 (qHsaCIP0030549)* and *B2M (qHsaCIP0029872)* and the SsoAdvanced™ Universal Probes Supermix (BioRad, 1725280) on a CFX96 Real-Time PCR System (BioRad). The mean of *HPRT* and *B2M* Cts was used for normalization. Relative mRNA levels were transformed into linear form by the 2 (-ΔΔCt) method (Bookout and Mangelsdorf, 2003). Fold changes of RNA expression (RQ) relative to unstimulated controls were used for statistical analysis.

The fibroblast and keratinocytes were lysed and RNA isolated by the RNeasy Midi Kit (#75144, Qiagen, Germany) according to the manufacturer’s instructions. RNA concentration was measured with a Nanodrop One instrument (Thermo Scientific) with 340 nm baseline correction. cDNA was generated from 1 µg total RNA using Taqman Reverse Transcription Reagents (#N8080234, Thermo Fisher Scientific, MA, USA) with random hexamers according to the manufacturer’s protocol for mono-cultured cells, and with LunaScript RT SuperMix kit (E3010, New England Biolabs) for co-cultured cells. RT-qPCR analysis was carried out using TaqMan Gene expression Master Mix (#4369510, Life Technologies, Thermo Fisher Scientific) for mono-cultured cells, and with Luna Universal Probe qPCR MasterMix (M3004, New England Biolabs) for co-cultured cells. All RT-qPCR analysis were performed on QuantStudio5 (Applied Biosystems, Foster City, CA, USA) PCR System. The amplification protocol using TaqMan Gene expression Master Mix was initiated at 50°C for 2 min, followed by denaturation at 95°C for 10 min, then 40 cycles of denaturation at 95°C for 15 secs followed by annealing of TaqMan probes and amplification at 60°C for 1 min. The amplification protocol using Luna Universal Probe Master Mix was initiated with initial denaturation at 95°C for 1 min, followed by 43 cycles with denaturation at 95°C for 15 sec, followed by annealing and extension of TaqMan probes 60°C for 35 sec using the Fast cycling profile. *EEF1A1* was used for normalization, and the relative gene expression (RQ) was calculated against the unstimulated control by the comparative 2^-ΔΔCt^ method. RT-qPCR analysis was performed on 3 biological replicates, each with 3 technical replicates. All commercial TaqMan® primers (Thermofisher) and probes are listed:

Hs00265885_g1 (*EEF1A1*)

Hs00230071_m1 (*NFKBIZ*)

Hs00161488_m1 (*S100A7*)

**Cell culture**

Buffy coats were obtained from the blood bank Ålesund (Helse Møre og Romsdal, Norway) with approval by the Regional Ethics Committee (REK, ref.: 230804). Peripheral blood mononuclear cells (PBMC) were isolated from buffy coats by Lymphoprep^TM^ (AXIS-SHIELD PoC AS,1114547) density gradient and monocytes were selected by plastic adherence and differentiated into monocyte-derived macrophages (MDM) by culture in AIM V medium (Gibco, 12055083) with 5 % CTS™ Immune Cell Serum Replacement (ICSR) and 10 ng/mL M-Csf (Miltenyi Biotec, 130-093-963) for 5 d, followed by incubation without M-Csf. To induce inflammatory signaling, cells were treated with 1 µg/mL lipopolysaccharide (LPS, Sigma-Aldrich, L2630) and 50 ng/mL IFN-γ (Miltenyi Biotec, 130-096-481).

For T-cell isolation, PBMC obtained by density gradient were stored overnight in AIM V with 5 % ICSR and CD4+ T-cells were isolated by negative magnetic bead separation using the CD4^+^ T Cell Isolation Kit (Miltenyi Biotec, 130-096-533). T-cell purity was checked by staining for CD4 (Miltenyi, 130-113-790), CD3 (Miltenyi, 130-113-700) and CD14 (Miltenyi, 130-110-577) and analysis on a BD Accuri Flow cytometer (NTNU, Ålesund) with visualization in FCSalyzer 0.9.22-alpha and was routinely above 95 %. T-cells were activated by CD3/CD28 Dynabeads™, at 1 bead/2 cells (Gibco, 11161D) and cytokines (IL-1β (20 ng/mL, Invitrogen, A42508), PGE_2_ (5 µM, Cayman chemicals, 14010), IL-23 (25 ng/mL, Miltenyi Biotec, 130-095-757) for 3 d. For co-culture of MDMs with autologous T-cells^32^, T-cells were either kept in culture supplemented with 1 ng/mL IL-7 or were frozen until differentiation of MDMs. MDMs were pretreated with HRO or not, activated by LPS for 24 h. At the same time, T-cells were pretreated with HRO or not and finally added in fresh medium 1:1 on the activated MDMs, with or without co-treatment with HRO in the co-culture for 3 d.

THP1-Dual™ NF-κB-SEAP IRF-Luc Reporter monocytes (InvivoGen, thpd-nfis) were grown at a concentration of 0.5 - 2 mio cells/mL in RPMI 1640 (Thermofisher, 61870036) with 10 % heat-inactivated fetal bovine serum (FBS) (Sigma-Aldrich, F0804), 100 U/mL penicillin + 0.1 mg/mL streptomycin (Sigma-Aldrich, P4333) and 100 μg/mL Normocin™ (InvivoGen, ant-nr-1), according to the manufacturer’s instructions. To keep selection pressure, 10 μg/mL Blasticidin (InvivoGen, ant-bl-1) and 100 μg/mL Zeocin™ (InvivoGen, ant-zn-1) were added every second passage. For experiments, the cells were differentiated to macrophages by seeding with 40 ng/mL Phorbol 12-myristate 13-acetate (PMA; Sigma-Aldrich, P8139) and incubation for 3 d, followed by 1.5 d rest in fresh medium without PMA. To induce inflammatory signaling, cells were treated with 10 ng/mL LPS.

Human primary dermal fibroblasts (ATCC PCS-201-012) and human keratinocytes HaCaT (CLS Cell line service GmbH, cat no 300493) were cultured in Dulbecco´s modified Eagle´s medium (DMEM) high glucose supplemented with 10 % FBS, 100 U/mL penicillin and 100 µg/mL streptomycin (all purchased from Thermo Fisher Scientific) before seeding at passages 4-6 in the different co-culture experiments. For the co-culture experiments in Transwell® (TW) 12-well plates (Corning Costar Transwell, VWR) 1x10^5^ fibroblasts were seeded at the plastic well bottom and cultured for 2 d before 5.5x10^5^ keratinocytes were seeded on the TW filters and further co-cultured for 5 d before stimulation. Stimulation was performed for 4 d with or without 100 ng/mL IL-17A (Sigma-Aldrich, cat.no SRP0675) and with or without 5 or 50 µg/mL HRO (in same culture media and incubation conditions as described above), before sampling of media and cells for further analysis. For direct co-culture, 3.6x10^3^, 3 separately grown populations of fibroblasts were plated in each well of a 12-well plate for RNA and 9.864 x10^3^ cells in each well of 6-well plates for protein. After 2 d of cultivation, 1.5x 10^4^ , 3 separately grown populations of keratinocytes were added to each well of the fibroblast containing 12-well plates, and 4.11 x10^4^ cells to each well of the 6-well plates. Cells were co-cultured for 5 d before stimulation as in the TW system for 4 d. For proliferation studies, 2.0x10^3^ fibroblasts or 4.0x10^3^ keratinocytes were plated per well in a 96-well plate for monoculture monitoring. For co-culture monitoring, 500 fibroblasts were mixed with 2.5x10^3^ keratinocytes in a 96-well plate. Stimulant conditions were added to the cells the day after plating, and real-time cell confluence was subsequently monitored every 4 h using an IncuCyte® S3 Live-Cell Analysis System (Sartorius AG, Germany). All cells were maintained in 5 % CO_2_ at 37˚C.

#### **Bio-Plex**

For cytokine analysis from fibroblast and keratinocyte cell media, Bio-Plex Pro Human Cytokine 8-plex Assay (Bio-Rad, M50000007A) was used, and run on a Bio-Plex Multiplex immunoassay system (Bio-Rad, BioPlex® 200).

**Western blot**

Cells grown for protein analysis were washed twice with cold PBS before lysis in Pierce™ RIPA buffer (89900, Thermo Scientific) containing Protease and Phosphatase inhibitor (A32959, Thermo Scientific) for 30 min on ice. The lysates were subsequently centrifuged at 13,000 rpm for 20 min at 4°C. Protein concentrations were determined using DC Protein Assay (BioRad). 20 µg protein was loaded per well of SDS-PAGE NuPage 12 % Bis-Tris gels (NP0342BOX, Invitrogen), and gels were run in 3-(N-morpholino) propane sulfonic acid (MOPS) SDS running buffer (NP0001, Invitrogen) at 200 V for approximately 50 min. Following electrophoresis, the proteins were transferred onto nitrocellulose membranes (iBlot3 Transfer stacks, IB33001, Invitrogen) using an iBlot Gel Transfer Device (Invitrogen). All membranes were blocked with 2 % ECL Prime blocking agent (RPN418V, Cytiva) in TBS-tween for 1 h at room temperature (RT). The primary and secondary antibodies were diluted in a 0.2 % blocking agent and incubated for 1.5 h at RT (or overnight at 4 °C) with gentle shaking. Membranes were washed 3 x 10 min with TBS-tween after both incubations. ECL plex™ Rainbow™ Fluorescent marker from Cytiva (#RPN850E, MA, USA) was used as a molecular weight marker. Proteins were scanned and visualized using G: BOX Chemi XX6/XX9-(Syngene, India). Primary antibodies used; mouse anti-α-tubulin (1:10.000, #T5168, Sigma Aldrich, USA), mouse anti-Psoriasin (1:1000, NB100-56559, Novus Biological). Secondary antibody used; ECL Plex goat-a-mouse IgG Cy3 (cat.no PA43009, Cytiva).
